# Oleic acid differentially affects *de novo* lipogenesis in adipocytes and hepatocytes

**DOI:** 10.1101/2023.10.04.560581

**Authors:** Hannah B. Castillo, Sydney O. Shuster, Lydia H. Tarekegn, Caitlin M. Davis

## Abstract

Lipogenesis is a vital but often dysregulated metabolic pathway. We report super-resolution multiplexed vibrational imaging of lipogenesis rates and pathways using isotopically labelled oleic acid and glucose as probes in live adipocytes and hepatocytes. These findings suggest oleic acid inhibits *de novo* lipogenesis (DNL), but not total lipogenesis, in hepatocytes. No significant effect is seen in adipocytes. These differential effects may be due to alternate regulation of DNL between cell types and could help explain the complicated role oleic acid plays in metabolism.

---

Liver diseases have become a significant health concern worldwide, especially with both obesity and alcohol consumption on the rise.^1^ Non-alcoholic fatty liver disease (NAFLD) is the most common liver disease at a staggering 25% lifetime prevalence worldwide.^2^ Liver cancer is among the top five leading causes of cancer-related deaths and is projected to surpass one million global cases by 2025.^3^ Dysregulation of *de novo* lipogenesis (DNL) is a significant factor in these diseases.^4^ While several inhibitors have been identified and developed,^5^ it is unlikely that a single enzyme holds the key to mediating these diseases. Rather, it is important to study control of the DNL pathway at the cellular level, including how responses to external stimuli, such as scavenging pathways, adjust overall rates of DNL and total lipogenesis. Thus, observing competing lipogenic pathways *in cellulo* is crucial to answering these pressing questions.

As a monounsaturated fatty acid, dietary oleic acid (OA) enters lipogenesis during fatty acid synthesis and undergoes esterification into lipid droplets (LDs, Fig. 1A).^6^ While fatty acid overload is typically associated with hepatic steatosis and the onset of diseases such as obesity and NAFLD, counterevidence suggests that unsaturated fatty acids act as a protectant from cell death.^6–8^ Oleic acid in particular has been shown to prevent cholesterol synthesis and liver lipotoxicity,^6^ while simultaneously affecting LD morphology and formation in ways characteristic of disease (increasing both their number and size).^9,10^ This suggests OA can specifically regulate deleterious aspects of lipogenesis associated with increased general lipid content.^8^ Enzymatic activity assays and mRNA studies have indicated OA may regulate cholesterol generation by inhibiting the expression and activity of acetyl-CoA carboxylase (ACC, Fig. 1A), at least short term (<4 h).^7,11^ Additionally, unsaturated fatty acids like OA have been shown to decrease expression of and inhibit sterol regulatory element binding protein 1c (SREBP-1c), a master regulator of DNL.^12–14^ Thus, a better understanding of morphologic and metabolic effects of OA on DNL and other lipid pathways is needed to elucidate its role in disease.

**Figure 1.**
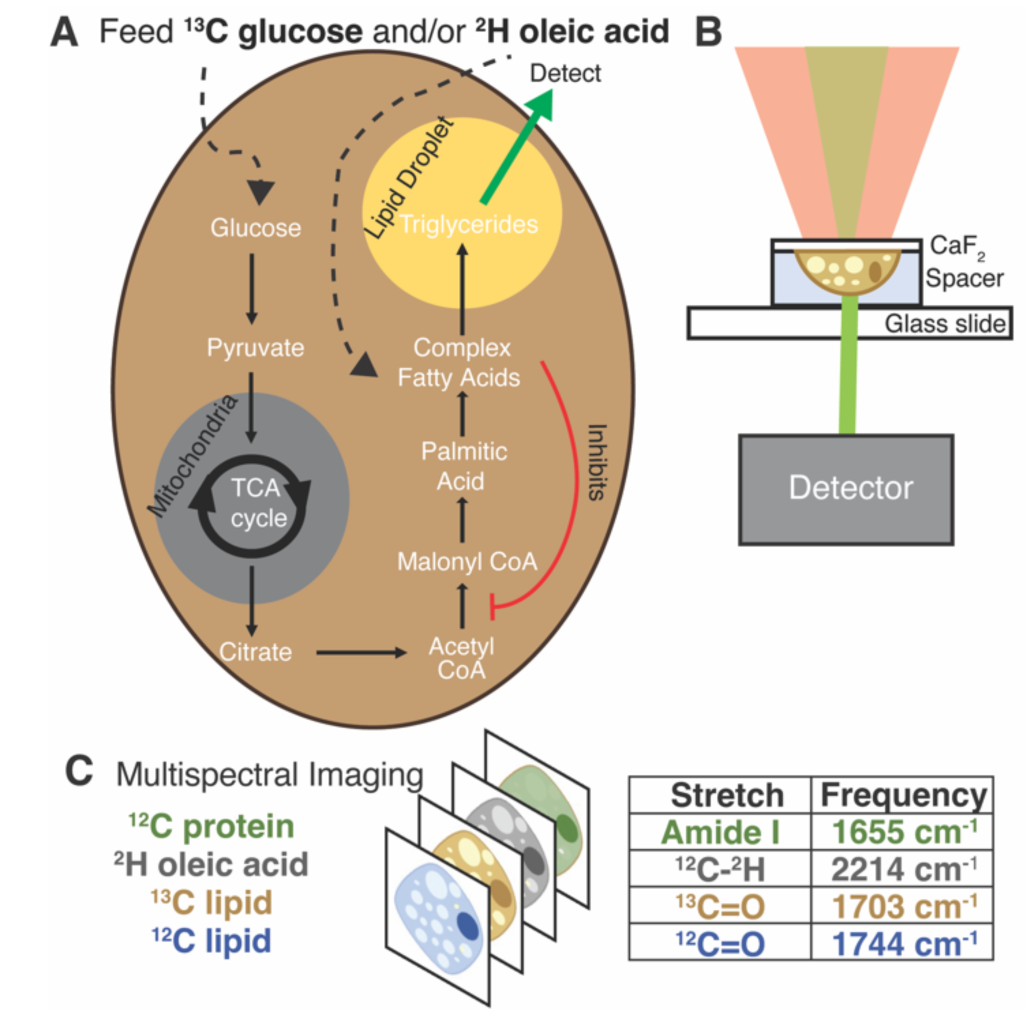
Schematic of experiments and data collection. (A) Isotope labelling of *de novo* lipogenesis and fatty-acid scavenging pathways. (B) Live cell data collection by OPTIR. Cells grown on a calcium fluoride cover slip are mounted in PBS using a spacer. (C) Multispectral imaging at single frequencies of interest.

Measuring metabolic pathways directly *in vivo* would provide this vital information, however, techniques for doing so are limited. Fluorescently labelled lipogenesis precursors like glucose and fatty acids are non-metabolizable or significantly disruptive. Other glucose analogs present a similar issue and are primarily used in uptake assays.^15^ The exception is isotope labelling of precursors, which is nonperturbative, nontoxic, and well metabolized.^15–17^ While isotopic labels are easy to incorporate and have been used extensively for mass spectrometry experiments,^18,19^ a non-destructive technique is necessary to monitor these probes in living cells. Furthermore, research has shown that metabolism is heterogeneous throughout the cell, with external stressors stimulating condensation of metabolic components and other regulating factors.^20^ This presents a need for techniques that can provide both spatially and temporally resolved metabolic information.

Vibrational microspectroscopy is a promising means of tracking metabolism throughout the cell using nonperturbative isotopic probes.^15–17,21^ Biomolecules have unique and specific vibrational signatures and can provide information about the surrounding environment.^22^ FTIR and Raman microspectroscopy have been successfully used in cellular studies, but each present its own challenges when working with live cells.^23,24^ Raman-based imaging is restricted by the small Raman scattering cross section, resulting in long imaging times, high powered lasers, and limited spectral coverage, making live cell work difficult.^24^ FTIR is limited by strong IR absorbance of water overlapping with the peaks of many other biomolecules and spatial resolution in the micron range, which is larger than most cellular organelles and therefore does not provide sub-cellular information.^16,25^

Our approach, optical photothermal infrared microscopy (OPTIR), addresses the issues posed by traditional Raman and FTIR imaging techniques. OPTIR is a nonperturbative and label-free super-resolution infrared imaging method. In this pump-probe system the sample is “pumped” with an IR beam, inducing a photothermal effect, which is monitored by a change in intensity of the probe, a visible laser. This is then amplified and reconstructed into an IR spectrum. The diffraction limit is based on the probe laser, bringing the spatial resolution below 500 nm. The specific geometry also leads to significantly less signal contribution from water.^17,26^ OPTIR has previously been used on fixed samples to obtain information on metabolic rates and chemical compositions,^27,28^ but only recently has this technique been applied to live cell imaging.^17^

We previously used OPTIR to track rates of DNL by feeding ^13^C glucose to differentiated 3T3-L1 cells (adipocytes) and monitoring incorporation of the labelled carbon into the ester carbonyl of triglycerides in both live and fixed cells.^17^ Here, we extend this study to the hepatocyte-derived Huh-7 cancer cell line (hereafter referred to as hepatocytes) because hepatic DNL is especially relevant to NAFLD and other metabolic diseases. Further, we incorporate a multiplexed imaging approach to monitor DNL and fatty-acid scavenging concomitantly (Fig. 1). Multiplexed imaging is widely used in single-cell studies because it provides a more comprehensive view of intertwined pathways.^23,29^

Multiplexed imaging in both live hepatocytes (Fig. 2A) and adipocytes (Fig. 2B) can be achieved because the vibrational probes used exhibit strong IR bands easily distinguishable amongst the fingerprint of other biomolecules (Fig. 2C). Briefly, by feeding Huh-7 and differentiated 3T3-L1 cells both ^2^H oleic acid and ^13^C glucose, the regulatory effect of oleic acid and resulting competing scavenging pathways can be observed (Fig. 1A). Live cell imaging is achieved by growing cells on a CaF_2_ coverslip and mounting it in buffer on a glass slide (Fig. 1B). Rates of DNL and OA scavenging/esterification are determined by multispectral imaging (Fig. 1C), which are collected sequentially as single wavenumber images set to frequencies associated with metabolism of the isotope-labelled probes as well as the protein amide-I and unlabelled lipid carbonyl stretch.

**Figure 2.**
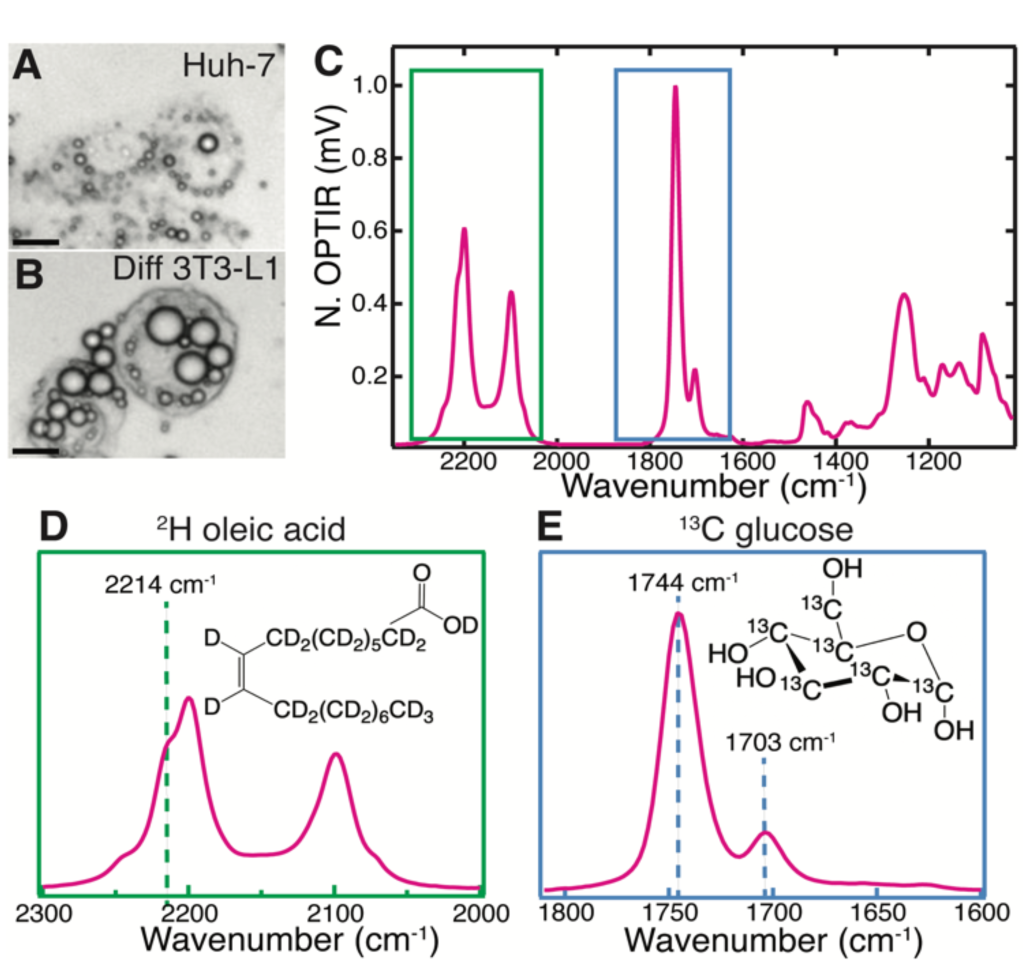
Live cell characterization of the cell lines (A) Huh-7 and (B) differentiated 3T3-L1 after feeding with ^2^H oleic acid and ^13^C glucose. Scale bars are 20 μm. (C) Full IR spectra of a point within a Huh-7 cell LD 48 hours after feeding ^2^H OA. (D) ^2^H-C from the ^2^H oleic acid (structure in inset) which has been esterified in LDs. 2214 cm^-1^ is assigned to ^2^H-C stretching. (E) ^13^C=O stretch of triglycerides at 1703 cm^-1^ where the ^13^C comes from metabolized ^13^C glucose (structure in inset). Pictured alongside the unlabelled ^12^C=O band at 1744 cm^-1^.

The full spectrum of ^2^H oleic acid (Fig. S1A) shows strong peaks at 2200 cm^-1^ and 2100 cm^-1^ attributed to C-^2^H stretches in the “cell silent region”.^16^ These peaks also appear in LDs of cells fed ^2^H OA (Fig. 2D, Fig. S1B). Shoulders at 2214 cm^-1^ and 2246 cm^-1^ are assigned to C-^2^H stretches of the singular C=C-^2^H in OA.^30,31^ The 2214 cm^-1^ shoulder provides a clear frequency from which to collect single wavenumber images to specifically measure uptake and esterification of ^2^H OA. Free OA is generally not visible outside of LDs because the concentrations are below the detection limit of the instrument. The OA is esterified into triglycerides before storage in LDs, but this does not cause shifts in the vibrational bands of the C-^2^H stretches.^32^

In the case of ^13^C glucose, as the glucose is metabolized the ^13^C is eventually incorporated into triglycerides in LDs via DNL (Fig. 1A). An increase in reduced mass affects the vibrational frequency, resulting in a redshift of lipid vibrational bands.^33^ The most significant is the redshift of the triglyceride ^12^C=O ester carbonyl stretch (hereafter referred to as ^12^C lipid band) from 1744 cm^-1^ to 1703 cm^-1^, which we use to monitor ^13^C incorporation (Fig. 2E).^17^

To ensure that the technique used previously on adipocytes^17^ can be applied to hepatocytes, a similar experiment was performed in which Huh-7 cells were incubated over the course of 72 hours with ^13^C glucose as sole source of glucose and imaged live. As anticipated, the resulting images depict smaller LDs compared to those of adipocytes, but similar ^13^C=O/^12^C=O lipid ester carbonyl ratios and therefore rates of DNL (Fig. S2). In this work, both live and fixed cell data was collected using multispectral single wavenumber imaging as it is approximately 25 times faster than full hyperspectral imaging. ^17^ This enables increased statistics with similar, though not as detailed, spectroscopic information.

Live cell multispectral imaging was then performed on Huh-7 (Fig. 3) and differentiated 3T3-L1 cells (Fig. S3) fed both ^2^H OA and ^13^C glucose. Trends were confirmed with fixed cell multispectral imaging (Fig. S4-5). We found that both cell lines readily took up ^2^H OA and sequestered it in LDs by 24 hours. Ratio images of the ^2^H lipid band of ^2^H OA to the ^12^C lipid band reveal that ^2^H OA concentrations remain constant in LDs over the 72 hour time period (Fig. 3B, Table S1). Ratio images of the ^13^C lipid band to the ^12^C lipid band specifically track DNL because the ^13^C in the red-shifted lipid band comes only from the ^13^C glucose. Figure 3C shows the increasing ratio of ^13^C labelled lipid, allowing us to estimate the rate of DNL (Fig. 3, Table S1-4). Notably, ^2^H OA and ^13^C glucose signal was primarily confined to LDs in living cells (Fig. 3A-D, Fig. S3B-D). 3T3-L1 cells had a much lower ratio of ^2^H lipid to total lipid in both live and fixed cells, but similarly showed a constant ratio from 24 to 72 hours (Fig. S3, Table S5-8). This may be due to higher lipid content and larger LDs in adipocytes as compared to hepatocytes, leading to a lower ratio of OA to total lipid. In agreement with prior work,^17^ there was significant cell to cell variability in both cell lines as well as limited heterogeneity of ratios within cells.

**Figure 3.**
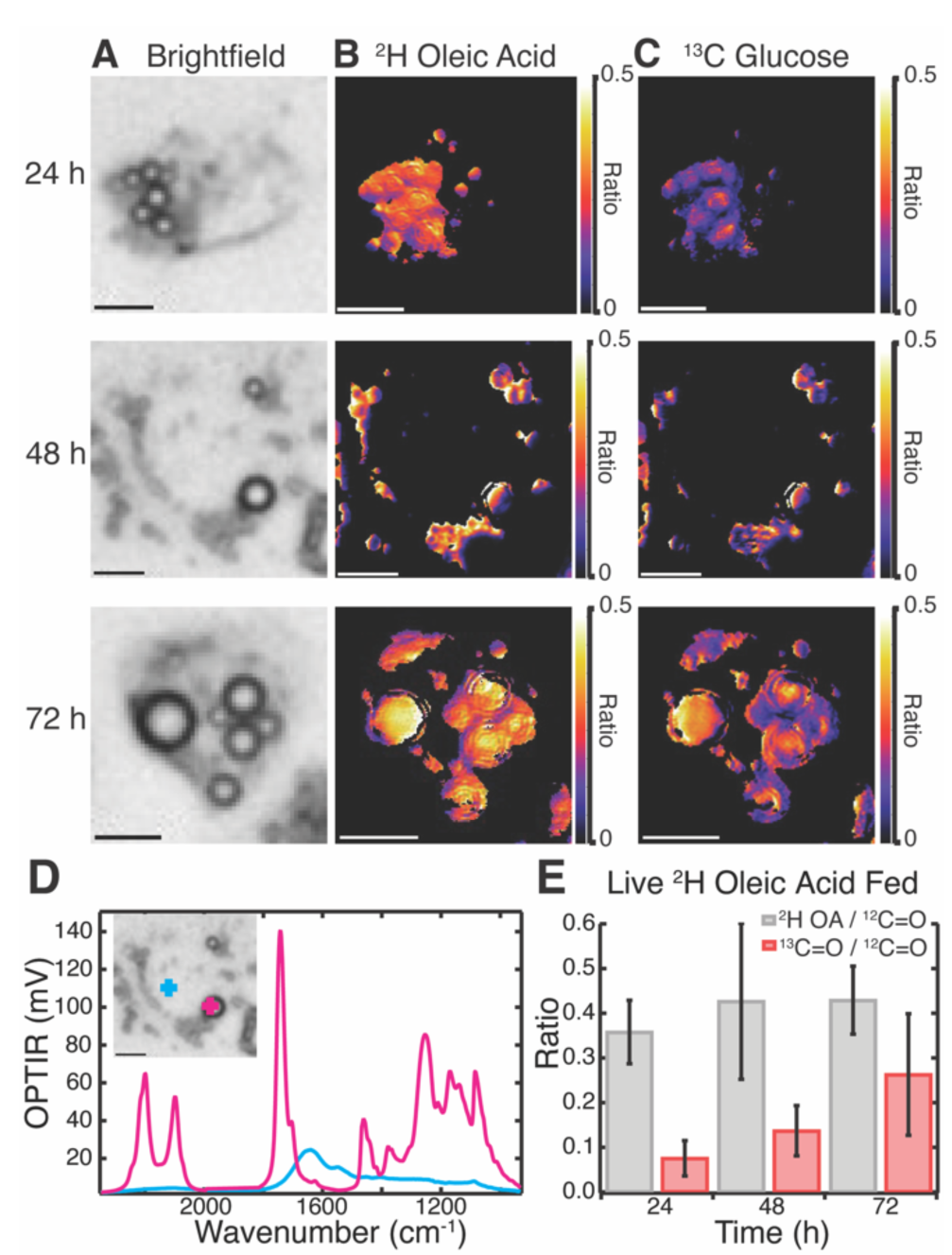
Visualization of rates of *de novo* lipogenesis in live Huh-7 cells fed with both ^2^H OA and ^13^C glucose. (A) Brightfield images of live cells. (B) ^2^H OA ratio images of ^2^H lipid to the ^12^C lipid band, after correction. (C) ^13^C glucose ratio images of the ^13^C lipid band to the ^12^C lipid band, after correction. (D) Representative spectra of the live Huh-7 cell at the 48-hour time point both inside a LD (pink) and outside the droplet (blue) (E) Average ratios of ^2^H lipid band (grey) and ^13^C lipid band (red) to ^12^C lipid band across many cells (N = 14). Scale bars are 10 μm.

Huh-7 cells showed a marked increase in both number and size of LDs after ^2^H OA feeding (Fig. S6), agreeing with previous literature.^10,34^ At the 48 hr time point, there were on average 9.2 ± 4.7 LDs in OA fed cells compared with 5.6 ± 2.4 LDs in vehicle fed (BSA only) conditions (Table S9). Similarly, LD diameter was 2.5 ± 0.8 μm in ^2^H OA fed conditions and 2.1 ± 0.5 μm in vehicle fed conditions (Table S9). The differences in both number and diameter of LDs between the two feed conditions were both extremely significant with p values below 0.0005.

Interestingly, ^2^H OA fed 3T3-L1 cells had much higher levels of DNL as compared to ^2^H OA fed Huh-7 cells with a final ^13^C/^12^C lipid ratio of ∼0.7 as compared to ∼0.3 (Fig. S3E and Fig. 3E red), respectively. To further investigate this, we compared the average ratios of ^2^H OA fed cells to cells fed only the BSA vehicle without OA (Fig. 4). Both sets contained reduced FBS to limit other sources of fatty acid scavenging. In Huh-7 hepatocytes, at every timepoint the ratio of ^13^C lipid to ^12^C lipid was dramatically and significantly (p < 0.05) reduced in ^2^H OA fed conditions (Fig. 4A). Conversely, the ratio remained largely unaffected in 3T3-L1 adipocytes (Fig. 4B). The same relationship was observed in fixed Huh-7 and 3T3-L1 cells (Fig. S7, Table S1-8). Therefore, DNL is downregulated in ^2^H OA fed Huh-7 cells and unchanged in ^2^H OA fed 3T3-L1 cells.

**Figure 4.**
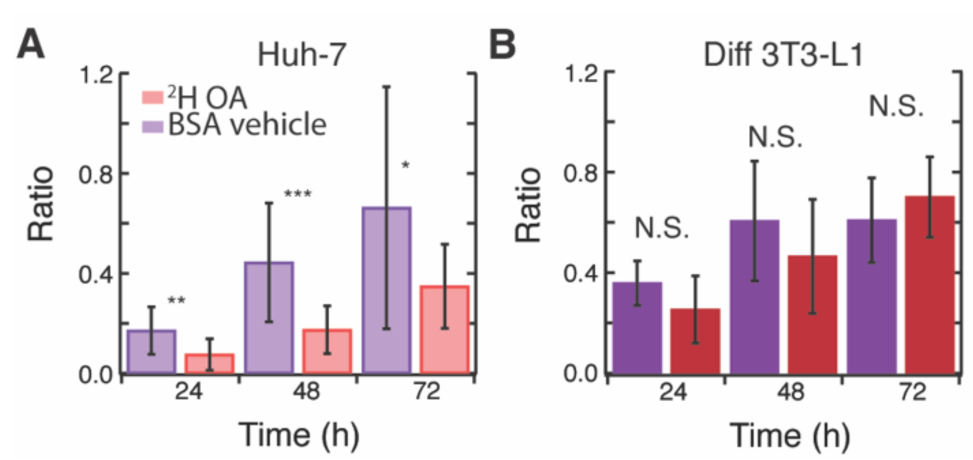
Effect of OA feeding on rates of *de novo* lipogenesis. Ratio of ^13^C lipid to ^12^C lipid in live (A) Huh-7 and (B) 3T3-L1 cells fed with both ^2^H oleic acid and ^13^C glucose (red) or BSA vehicle and ^13^C glucose (purple) at their respective time points. N = 8-16 cells per time point * p-value < 0.05 ** p-value < 0.005 ***p-value < 0.0005 N.S. p-value > 0.05

However, as evidenced by the increased number and size of LDs (Fig. S6, Table S9), the total ^12^C lipid content also increases in ^2^H OA fed Huh-7 cells. This increase in LDs has been previously reported^10,34^ and could artificially decrease the ratio used to approximate DNL. However, the rate of DNL is measured by the rate of change of the ^13^C/^12^C lipid and, therefore, should not be affected by a change in total lipid content. Thus, we fit the average ratio data of the Huh-7 cells and found very different rates with a linear slope of approximately 0.01 ^13^C/^12^C lipid ratio per hr for the BSA vehicle fed cells and 0.005 ^13^C/^12^C lipid ratio per hr for the ^2^H OA fed cells for both live and fixed cells (Fig. S8). DNL is slowed by half by OA feeding. In 3T3-L1 cells no difference is seen between the ratio in the BSA vehicle and ^2^H OA fed conditions at any time point (Fig. 4B).

We found that ^2^H OA feeding in Huh-7, but not differentiated 3T3-L1 cells, is correlated with both an increase in total lipogenesis and therefore LD size and number and a decrease in *de novo* lipogenesis. Reduced DNL after OA feeding has previously been reported in glioma cells and was associated with OA inhibition of ACC activity and expression.^7,35^ In line with our data, OA is also known to inhibit *de novo* fatty acid and cholesterol synthesis but increase triglyceride and general lipid content.^7,11^ OA has also been shown to be protective against cell death in hepatocytes, even when inducing steatosis.^36,37^ This may be due to its differential effects on DNL and total lipogenesis as DNL primarily generates palmitate, a fatty acid associated with cell death.^4,37^ Interestingly, OA feeding had little impact on adipocyte DNL, despite being readily taken up and stored in LDs (Fig. S3). The difference between the two cell lines could simply be a concentration effect as cultured adipocytes contain more lipid, perhaps making them less sensitive to small changes in the synthesis of new lipid. However, this seems unlikely as rates of DNL are similar between adipocytes and hepatocytes in the BSA vehicle fed conditions (Fig. 4). Alternatively, OA is known to reduce the activity of SREBP-1c,^12,13^ a regulator of DNL enzymes, including ACC, that is sufficient and necessary for DNL in hepatocytes.^38,39^ In adipocytes, however, SREBP-1c knockout does not significantly decrease expression of lipogenic enzymes^40^ and overall SREBP-1c seems to be a much smaller player for adipocyte DNL.^39,41^ We propose that OA may specifically regulate SREBP-1c and therefore hepatocyte ACC. This explains its dramatic effect on hepatocyte DNL but not adipocyte DNL. Further research is needed to fully disentangle the relationship between fatty acid scavenging and DNL, especially across cell lines. This work uniquely highlights the ability of OPTIR not only to observe multiplexed vibrational probes in living cells, but also provide insight on competing metabolic pathways.

This work was supported by National Institutes of Health (NIH) grant R35 GM151146, the Research Corporation for the Advancement of Science, and the Gordon and Betty Moore Foundation. H.B.C. and S.O.S. were partially supported by the NIH under Biophysics Training Grant No. T32 GM 008283 and S.O.S was partially supported by a National Science Foundation Graduate Research Fellowship under Grant No. DGE-2139841. L.H.T. was partially supported by the Yale College Dean’s Office through the STARS II Program.

