## Supplemental Methods and Figures for "Oleic acid differentially affects *de novo* lipogenesis in adipocytes and hepatocytes"

**Contents**

Experimental Methods

Materials

Huh-7 cell culture and ^13^C and ^2^H labeling

Differentiated 3t3-L1 cell culture and ^13^C and ^2^H labeling

Cell fixing

Live cell chamber construction

OPTIR data collection

Data analysis

Lipid droplet size analysis

Presentation of results and statistics

Supplemental Figures

Figure S1. Full assignments of ^2^H oleic acid spectrum and cell fed with probes

Figure S2. Live Huh-7 cell after 72 hours of feeding ^13^C glucose

Figure S3. Visualization of rates of DNL in live differentiated 3T3-L1

Figure S4. Visualization of rates of DNL in fixed Huh-7 cells

Figure S5. Visualization of rates of DNL in fixed differentiated 3T3-L1

Figure S6. Fixed cell rates of ^13^C incorporation in the vehicle and ^2^H oleic acid

Figure S7. Lipid droplet comparison brightfield images

Figure S8: Slopes of ^13^C incorporation rates

Supplemental Tables

Table S1: Average ratios of live and fixed Huh-7 cells fed with ^2^H OA and ^13^C glucose

Table S2: Average ratios of live and fixed Huh-7 cells fed with ^13^C glucose in BSA

Table S5: Average ratios of live and fixed adipocytes fed with ^2^H OA and ^13^C glucose

Table S6: Average ratios of live and fixed adipocytes fed with ^2^H OA in BSA

Note: Tables S3,S4,S7, S8, and S9 are found in the attached excel files.

References

Experimental Methods

**Materials**

Unless otherwise specified, all reagents were sourced from Sigma-Aldrich.

**Oleic Acid Conjugation Protocol**

For feeding, ^2^H oleic acid (Cambridge Isotope Laboratories, Tewksbury, MA) was bound to fatty acid-free bovine serum albumin (BSA) using a protocol from Shi *et al*.^1^ In a 2 mL clear glass container (VWR, Radnor, PA), Milli-Q purified water, 20 µL of 1 M NaOH, and ^2^H oleic acid were combined and heated in a 70 °C water bath until clear. A solution of 150 mg/mL fatty acid-free bovine serum albumin (BSA) was then added to the ^2^H oleic acid solution and heated again at 37 °C in a water bath until clear. The solution was then filtered with a 0.2 μm filter into another 2mL clear glass container. Oleic acid concentration was approximately 3 mM.

**Huh-7 cell culture and ^13^C and ^2^H labeling**

Huh-7 cells (gift from Lars Plate, Vanderbilt University) were cultured in 75 cm^3^ sterile vented cap tissue culture flasks (Corning, Corning, NY). Cells were grown to confluence in Dulbecco’s modified Eagle’s medium containing 4.5 g/L glucose and L-glutamine (DMEM, Corning) supplemented with 10% fetal bovine serum (FBS) and 1% penicillin/streptomycin (Thermo-Fischer, Waltham, MA) under standard conditions (37° C, humidified atmosphere, 5% CO_2_). At 80 % confluence, cells were trypsinized and replated on CaF_2_ coverslips (20 x 20 x 0.35 mm, Crystran, Poole, U.K.) in 35 mm diameter sterile Petri dishes (Corning) for live samples and CaF_2_ coverslips (10 x 10 x 0.35 mm) in a 24 well cell culture plate (Corning) for fixed samples. After a day of allowing cells to adhere to the coverslips, the media was replaced with glucose-free DMEM supplemented with 1% FBS, 1% penicillin/streptomycin, ^13^C glucose to 4.5 mg/mL, and either 2% BSA or ^2^H oleic acid conjugated with BSA.

**Differentiated 3T3-L1 cell culture and ^13^C and ^2^H labeling**

3T3-L1 cells (ATCC, Manassas, VA) were prepared as previously described.^2^ Briefly, low passage number (<10), pre-differentiated cells were grown to confluence in Dulbecco’s modified Eagle’s medium containing 4.5 g/L glucose and L-glutamine (DMEM, Corning) supplemented with 10% calf bovine serum (CBS, ATCC) and 1% penicillin/streptomycin (Thermo-Fischer) under standard conditions. Two to three days post confluence, media was changed to DMEM containing 10% fetal bovine serum (FBS, Corning), 1% penicillin/streptomycin, 20 μg/mL insulin, 250 nM dexamethasone, and 500 μM isobutylmethylxanthine. Two to three days after differentiation, media was changed to DMEM containing 10% fetal bovine serum (FBS, Corning), 1% penicillin/streptomycin and 20 μg/mL insulin. After 2-3 days, cells were trypsinized and replated on CaF_2_ coverslips (20 x 20 x 0.35 mm; Crystran.) in 50 mm dishes (Corning). Media was exchanged every 2-3 days until lipid droplets formed and stabilized (5-7 days). At this point, medium was changed to glucose-free DMEM supplemented with 1% FBS, 1% penicillin/streptomycin, ^13^C- labeled glucose to 4.5 mg/mL, and either 2% BSA or ^2^H oleic acid conjugated with BSA.

**Cell fixing**

For fixed cell samples, both cell lines were fixed at the desired time points with 4% paraformaldehyde (PFA) in PBS (Corning) for 20 minutes at room temperature and washed three times with PBS followed by three times with Milli-Q purified water and allowed to air-dry before analysis.

**Live cell chamber construction**

After feeding, live cell samples from both cell lines were washed once with PBS and mounted in PBS on a glass microscopy slide (VWR) with double-sided tape spacers (Nitto, San Diego, CA) with a 5 μm thickness to keep the cells hydrated and sealed and minimize compression of the cell.

**OPTIR data collection**

All imaging was performed on a mIRage-LS IR microscope (Photothermal Spectroscopy Corporation, Santa Barbara, CA) integrated with a four-module-pulsed quantum cascade laser (QCL) system (Daylight Solutions, San Diego, CA) with a tunable range from 932 cm^-1^ to 2348 cm^-1^. Brightfield optical images were collected using a low magnification 10X refractive objective with a working distance of 15 mm. Spectra and infrared images were collected in co-propagating mode using a 40X Cassegrain objective with a working distance of 8 mm. Fixed cell spectra and images were collected in standard (reflective) mode and live cell spectra and images were collected in transmission mode. Data was collected with an IR laser power of 20 % and a probe power in the range of 5-11 %. All spectra and images were collected using PTIR Studio 4.5. (Photothermal Spectroscopy Corporation).

**Data analysis**

Images were processed in Fiji (NIH, Bethesda, MD).^3^ Spectra were analyzed in IGOR Pro 9 (Wavemetrics, Portland, OR). Live and fixed cell ratio images were generated in Python 3.10 in Colab (Google, Mountain View, CA).^4^

Ratio images must be corrected to account for overlapping signals of other biomolecules. This is done through a correction value, which removes contribution from the water band and protein amide I. Correction values are calculated for each cell line after analyzing single cell spectra in which no lipid signal is present. The intensity at the wavenumber at which lipid signal is collected (either ^13^C=O, ^12^C=O, or ^12^C-^2^H) is divided by the intensity at the amide I (1655 cm^-1^) or water band (2050 cm^-1^). These values are collected per cell across all time points and averaged to create a correction value with which to subtract off overlapping amide-I and water intensity for lipid intensities.

Fixed cell ^13^C=O/^12^C=O ratio images were corrected for the amide-I intensity using equation 1:

$$\frac{{}^{\text{13}}\text{C}}{{}^{\text{12}}\text{C}}\text{ ester carbonyl intensity = }\frac{A_{{}^{13}{C=O}}-bA_{Amide I}}{A_{{}^{12}{C=O}}}$$

**Equation 1**

Where *A* is the signal intensity of the frequency of the indicated biomolecule. The ^12^C=O stretch is at 1744 cm^-1^ for Huh-7 cells and at 1747 cm^-1^ for adipocytes. For both cell lines, the ^13^C=O stretch is at 1703 cm^-1^. The variable *b* is the correction value for the amide-I intensity, with *b* being 0.34 for Huh-7 cells and 0.31 for adipocytes. There was no correction done for fixed ^2^H-C/^12^C=O ratio images. Fixed cell ratios were only performed on regions with sufficient ^12^C=O signal intensity, defined as regions with 15% of the maximum value of ^12^C=O signal for Huh-7 cells and 5% for adipocytes which visually corresponded to the lipid rich areas of the cells as determined by examining the single wavenumber images at 1744 cm^-1^ and the brightfield view of lipid droplets.

Live cell ^13^C=O/^12^C=O ratio images were corrected for the amide I and water band intensity using equation 2.

$$\frac{{}^{\text{13}}\text{C}}{{}^{\text{12}}\text{C}}\text{ ester carbonyl intensity = }\frac{A_{{}^{13}{C=O}}-cA_{Amide I}}{A_{{}^{12}{C=O}}-dA_{Amide I}}$$

**Equation 2**

Where *c* is the correction value for $A_{{}^{13}{C=O}}$ and is 0.61 for Huh-7 cells and 0.66 for adipocytes and *d* is the correction value for $A_{{}^{12}{C=O}}$and is 0.31 for Huh-7 cells and 0.28 for adipocytes. Live cell ^2^H/^12^C ratio images were corrected for the water band using equation 3.

$$\frac{{}^{\text{2}}\text{H}}{{}^{\text{12}}\text{C}}\text{ ester carbonyl intensity = }\frac{A_{{}^{12}{C-{}^{2}H}}-fA_{water band}}{A_{{}^{12}{C=O}}-dA_{Amide I}}$$

**Equation 3**

The ^12^C -^2^H is measure from a shoulder at 2214 cm^-1^ for Huh-7 cells and at 2212 cm^-1^ for adipocytes. The water band is measured from 2050 cm^-1^ for both cell lines as this does not overlap with other ^12^C -^2^H stretches. The variable *f* is the correction value for the water band and is 0.87 for Huh-7 cells and 0.85 for adipocytes and *d* is the correction value discussed above. Live cell ratios were also only performed on regions with sufficient ^12^C lipid ester carbonyl signal intensity, defined as regions with 15% of the maximum value of ^12^C lipid ester carbonyl signal for Huh-7 cells and 5 % for adipocytes.

**Lipid droplet size analysis**

Lipid droplet size and diameter were determined using Fiji on cells at 48 hours after feeding with ^13^C glucose and either ^2^H oleic acid or BSA. The number of lipid droplets per cell were counted and the size of each lipid droplet determined using the line tool and measuring the diameter. These values were averaged over a total of 30 cells for both ^2^H oleic acid fed and BSA fed trials.

**Presentation of results and statistics**

Calculations for averages and statistical analysis were performed in Microsoft Excel (Microsoft, Redmond, WA). Bar graphs were generated by averaging the average ratios of N cells (Tables S1-8) at each time point. Error bars represent one standard deviation of average of the averages of the cells at each time point including propagated error from the standard error of each cell. Two tailed student’s t-tests were performed in Excel to determine significance between time points in trials.

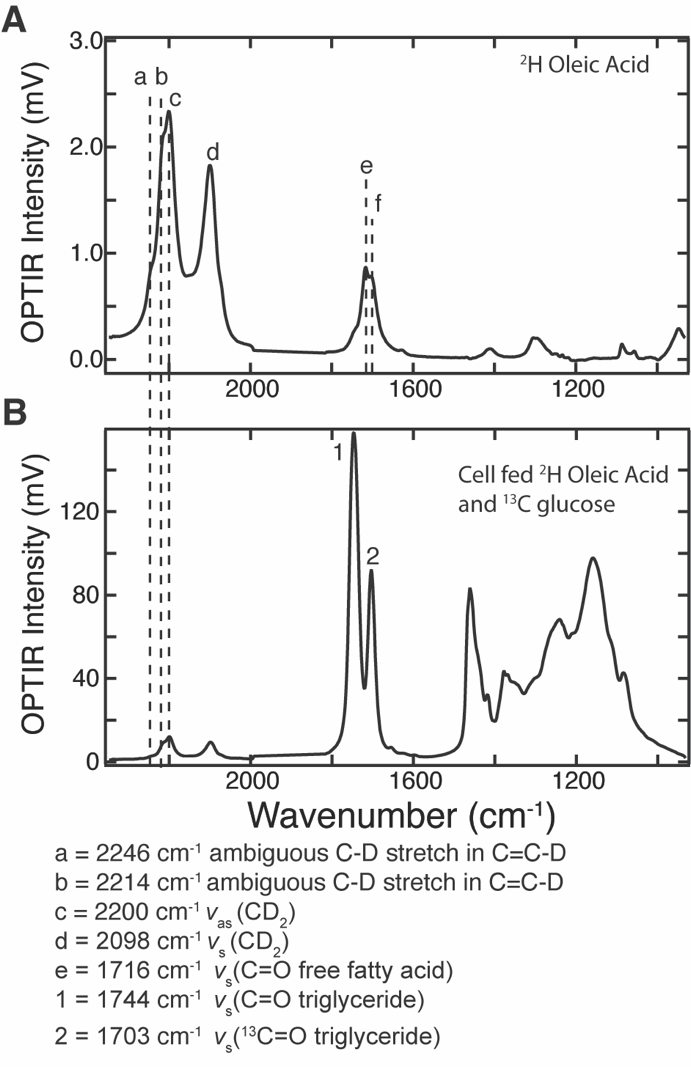

**Figure S1:** Full assignments of (A) ^2^H oleic acid IR spectrum^5,6^ and (B) a live differentiated 3T3-L1 cell 72 hours after feeding with ^2^H oleic acid and ^13^C glucose.

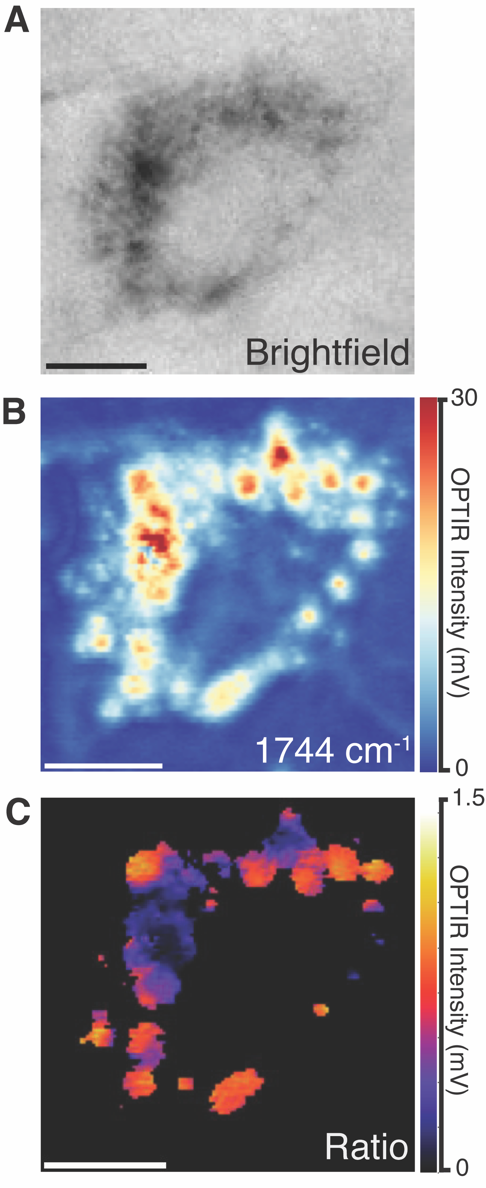

**Figure S2:** Rates of lipogenesis in live Huh-7 cells can be observed by OPTIR imaging approach previously used on adipocytes. (A) Brightfield image of a ^13^C glucose-fed Huh-7 cell at 72 hours. (B) Single wavenumber image collected at the ^12^C=O lipid band (1744 cm^-1^) and (C) the corresponding ratio image of ^13^C=O lipid ester carbonyl of triglycerides to ^12^C=O lipid band, after correction. Scale bars are 10 μm.

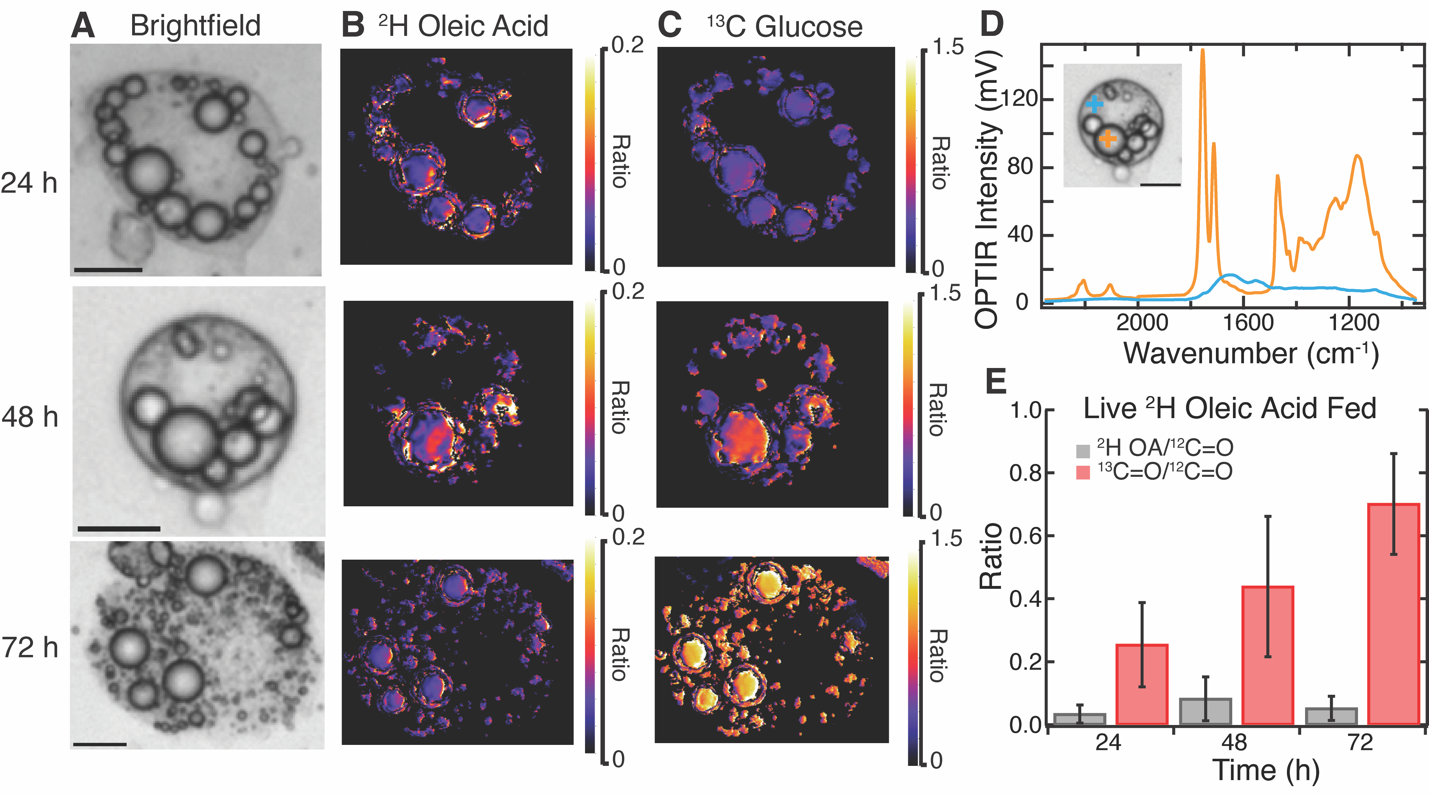

**Figure S3:** Visualization of rates of de novo lipogenesis in live differentiated 3T3-L1 cells fed with both deuterated oleic acid and ^13^C glucose for 24 to 72 h. (A) Brightfields images of live cells. B) ^2^H OA ratio images of ^2^H lipid to the ^12^C lipid band, after correction. (C) ^13^C glucose ratio images of the ^13^C lipid band to the ^12^C lipid band, after correction Scale bars are 10 μm. (D) A representative spectra of the live differentiated 3T3-L1 cell at the 48-hour time point. Orange trace was collected in a lipid droplet while blue was collected outside the droplet. (E) Average ratios of ^2^H lipid band (grey) and ^13^C lipid band (red) to ^12^C lipid band across many cells N=7-11 cells per time point.

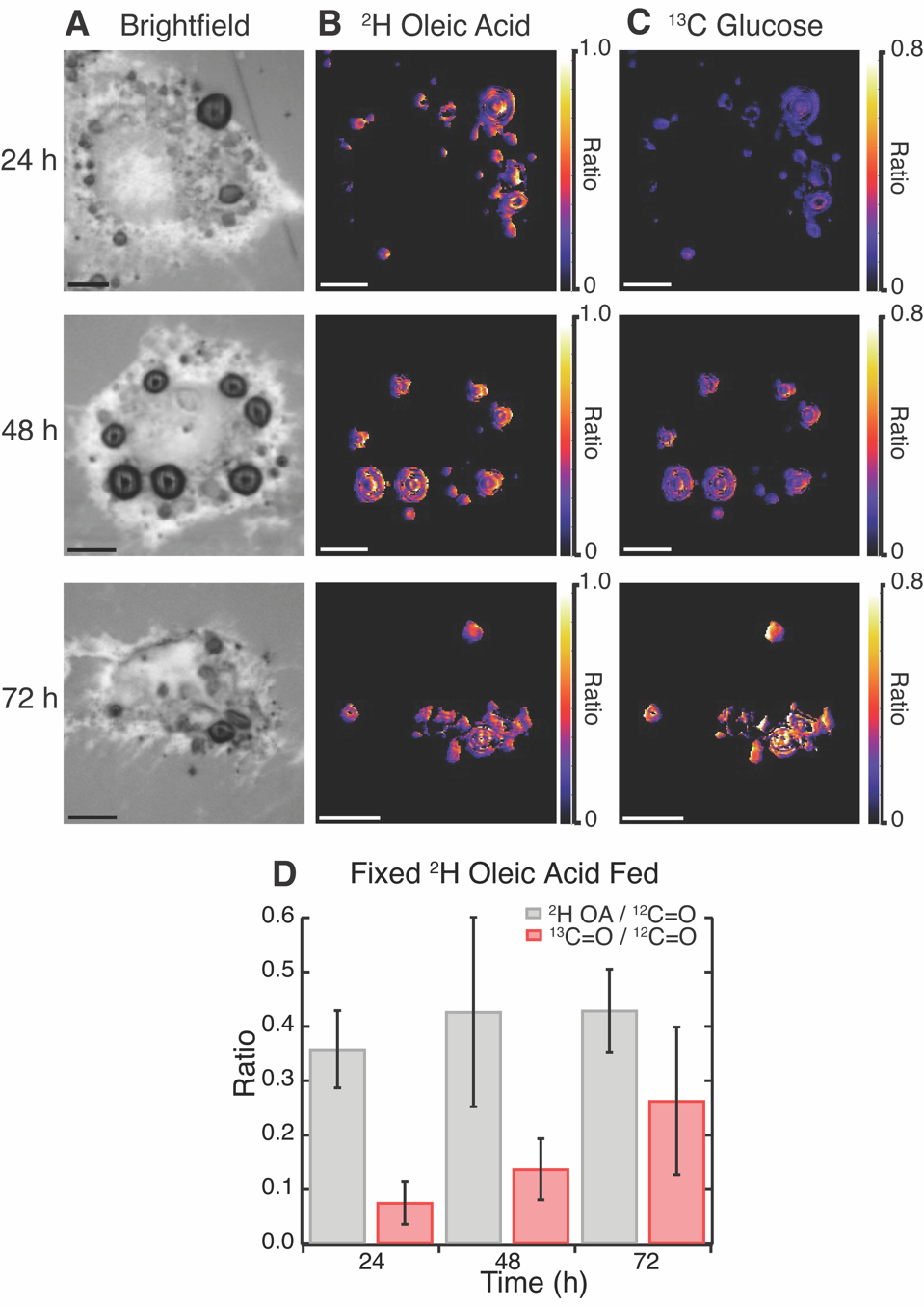

**Figure S4:** Visualization of rates of de novo lipogenesis in fixed Huh-7 cells after feeding of ^2^H oleic acid and ^13^C glucose for 24 – 72 h. (A) Brightfield images of fixed Huh-7 cells (B) ^2^H OA ratio images of ^2^H lipid to the ^12^C lipid band, after correction. (C) ^13^C glucose ratio images of the ^13^C lipid band to the ^12^C lipid band, after correction Scale bars are 10 µm. (D) Average ratios of ^2^H lipid band (grey) and ^13^C lipid band (red) to ^12^C lipid band across many cells. N=12-16 cells per time point.

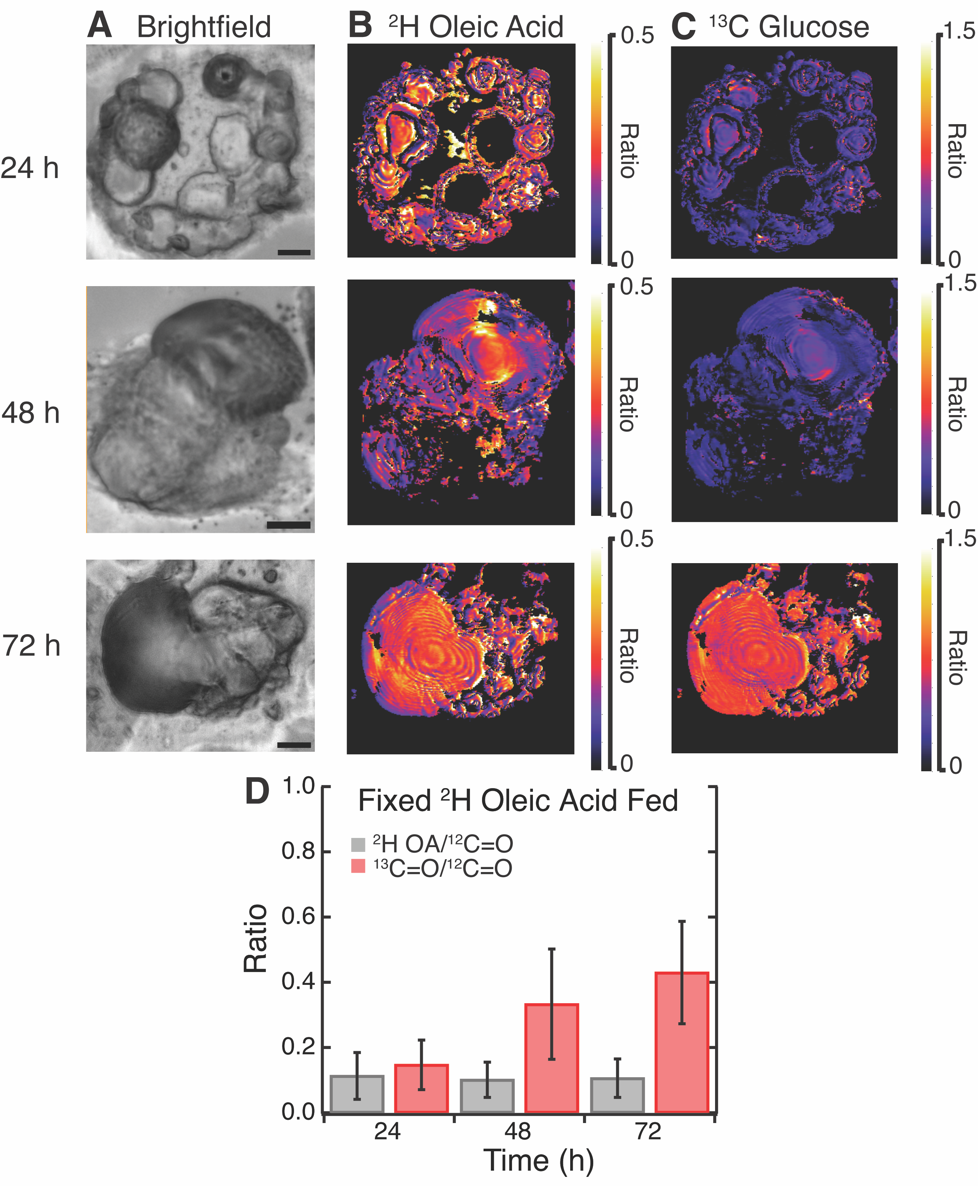

**Figure S5:** Visualization of rates of de novo lipogenesis in fixed differentiated 3T3-L1 cells after feeding of ^2^H oleic acid and ^13^C glucose for 24-72 h. (A) Brightfields of fixed differentiated 3T3-L1 cells. B) ^2^H OA ratio images of ^2^H lipid to the ^12^C lipid band, after correction. (C) ^13^C glucose ratio images of the ^13^C lipid band to the ^12^C lipid band, after correction Scale bars are 10 µm. (D) Average ratios of ^2^H lipid band (grey) and ^13^C lipid band (red) to ^12^C lipid band across many cells N=11-13 cells per time point

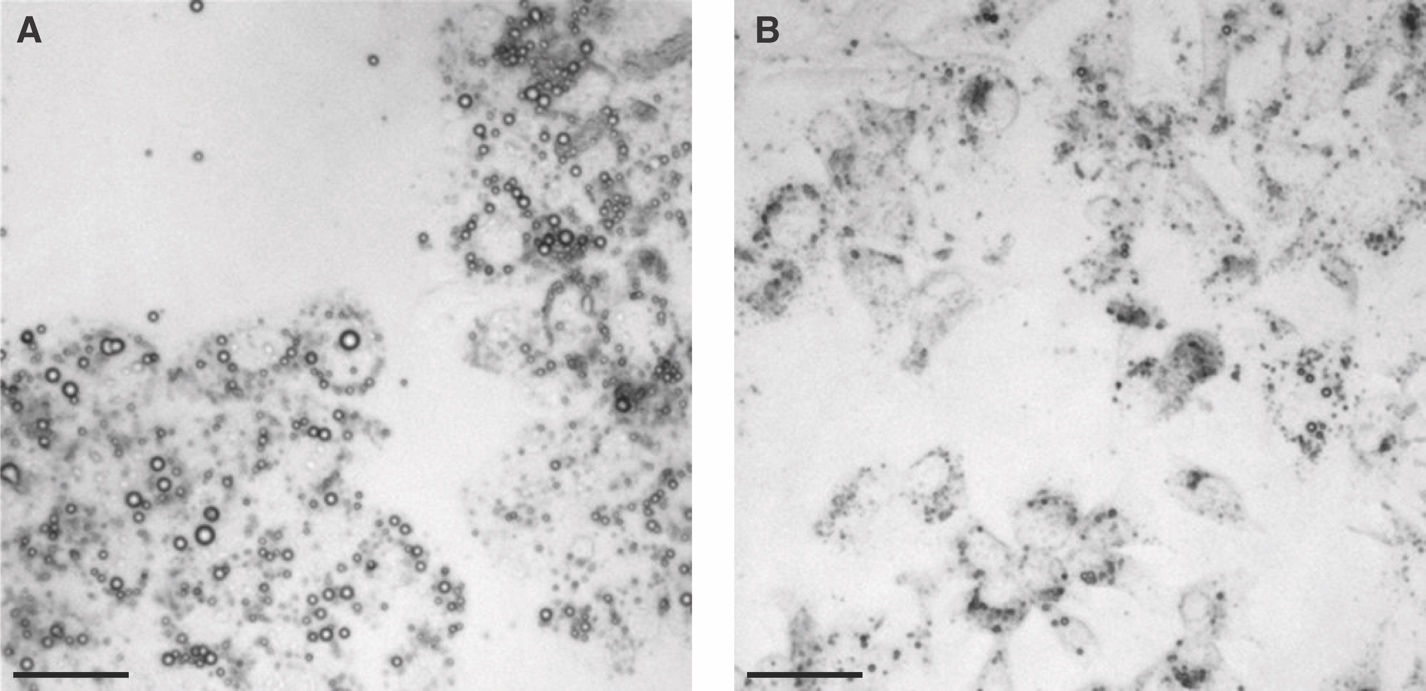

**Figure S6:** Lipid droplet comparison in Huh-7 cells at 48 hours after (A) feeding with ^2^H oleic acid and ^13^C glucose and (B) feeding with BSA and ^13^C glucose. Scale bars are 50 µm.

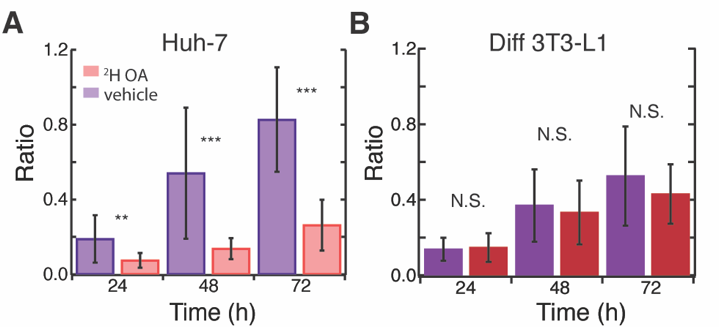

**Figure S7** Effect of OA feeding on DNL in fixed cells. Rates of uptake of ^13^C glucose in ^2^H OA trials (red) compared to in the BSA vehicle (purple) show that oleic acid has different effects between (A) fixed Huh-7 cells and (B) fixed differentiated 3T3-L1 cells. N= 7-16 cells per time point. * p-value< 0.05, ** p-value<0.005, ***p-value<0.0005

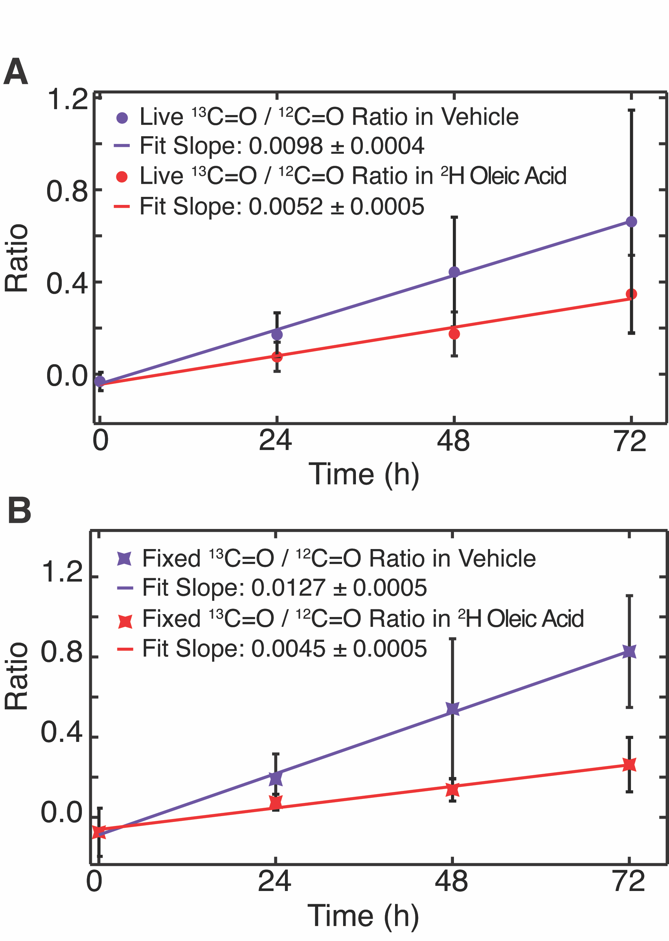

**Figure S8:** Slopes of the linear fit of ^13^C=O to ^12^C=O ratios in the vehicle and oleic acid at respective time points in (A) live and (B) fixed Huh-7 cells support the observation of a suppression of de novo lipogenesis with exposure to oleic acid.

Table S2: Average ratios of live and fixed Huh-7 cells fed with ^2^H OA and ^13^C glucose.

Table S3: Average ratios of live and fixed Huh-7 cells fed with ^13^C glucose in vehicle (BSA).

| *Live* |  |  |
| --- | --- | --- |
| Hour | N | Avg ^13^C=O/^12^C=O |
| 24 | 12 | 0.2 ± 0.1 |
| 48 | 12 | 0.4 ± 0.2 |
| 72 | 11 | 0.7 ± 0.5 |
| *Fixed* |  |  |
| Hour | N | Avg ^13^C=O/^12^C=O |
| 24 | 12 | 0.2 ± 0.1 |
| 48 | 16 | 0.5 ± 0.4 |
| 72 | 16 | 0.8 ± 0.3 |

| *Live* |  |  |  |
| --- | --- | --- | --- |
| Hour | N | Avg ^2^H OA /^12^C=O | Avg ^13^C=O/^12^C=O |
| 24 | 14 | 0.19 ± 0.07 | 0.08 ± 0.06 |
| 48 | 14 | 0.18 ± 0.08 | 0.17 ± 0.10 |
| 72 | 14 | 0.17 ± 0.07 | 0.4 ± 0.2 |
| *Fixed* |  |  |  |
| Hour | N | Avg ^2^H OA /^12^C=O | Avg ^13^C=O/^12^C=O |
| 24 | 12 | 0.36 ± 0.07 | 0.08 ± 0.04 |
| 48 | 16 | 0.4 ± 0.2 | 0.14 ± 0.06 |
| 72 | 16 | 0.39 ± 0.08 | 0.3 ± 0.1 |

Table S6: Average ratios of live and fixed differentiated 3T3-L1 cells fed with ^2^H OA and ^13^C glucose.

Table S7: Average ratios of live and fixed differentiated 3T3-L1 cells fed with ^13^C glucose in vehicle (BSA).

| *Live* |  |  |
| --- | --- | --- |
| Hour | N | Avg ^13^C=O/^12^C=O |
| 24 | 8 | 0.36 ± 0.09 |
| 48 | 8 | 0.6 ± 0.2 |
| 72 | 10 | 0.6 ± 0.2 |
| *Fixed* |  |  |
| Hour | N | Avg ^13^C=O/^12^C=O |
| 24 | 8 | 0.14 ± 0.06 |
| 48 | 9 | 0.4 ± 0.2 |
| 72 | 9 | 0.5 ± 0.3 |

| *Live* |  |  |  |
| --- | --- | --- | --- |
| Hour | N | Avg ^2^H OA /^12^C=O | Avg ^13^C=O/^12^C=O |
| 24 | 7 | 0.03 ± 0.03 | 0.3 ± 0.1 |
| 48 | 7 | 0.09 ± 0.07 | 0.5 ± 0.2 |
| 72 | 11 | 0.05 ± 0.04 | 0.7 ± 0.2 |
| *Fixed* |  |  |  |
| Hour | N | Avg ^2^H OA /^12^C=O | Avg ^13^C=O/^12^C=O |
| 24 | 11 | 0.11 ± 0.07 | 0.15 ± 0.08 |
| 48 | 11 | 0.10 ± 0.05 | 0.3 ± 0.2 |
| 72 | 13 | 0.11 ± 0.06 | 0.4 ± 0.2 |
